# Molecular characterization of mosquitoes (Diptera: Culicidae) from the tropical rainforest of Sierra Nevada de Santa Marta, Colombia

**DOI:** 10.1101/2020.06.04.135095

**Authors:** Andrew Muñoz-Gamba, Katherine Laiton-Donato, Erick Perdomo-Balaguera, José Usme-Ciro, Gabriel Parra-Henao

## Abstract

**BACKGROUND:** The Sierra Nevada de Santa Marta rainforest has diverse fauna due to its position in northern Colombia, a Caribbean region with predominantly tropical, dry, and rainforest ecosystems in which there is a high diversity of mosquito species that may act as arbovirus vectors.

**OBJECTIVES:** The present study reports the molecular characterization of select mosquito species in this rainforest.

**METHODS:** Manual capture methods were used to collect mosquitoes, and the specimens were identified via classical taxonomy. The *COI* marker was used for species confirmation, and phylogenetic analysis was performed, using the neighbor-joining method, with the Kimura-2-Parameters model.

**FINDINGS:** *Aedes serratus*, *Psorophora ferox*, *Johnbelkinia ulopus*, *Sabethes cyaneus*, *Wyeomyia aporonoma*, *Wyeomyia pseudopecten*, *Wyeomyia ulocoma* and *Wyeomyia luteoventralis* were identified and intra-species variation >2% for most species.

**MAIN CONCLUSIONS:** We report the first records on the genetic variability of mosquitoes in this area and phylogenetic reconstructions allowed for identification at the species level, and the corroboration by means of classical taxonomy suggested complementarity of both methods, which may be employed when morphological or molecular data are poor or not available. The genetic and morphological characterization of jungle mosquito populations will help to understand their biology.

## Introduction

The family Culicidae comprises 3,600 species, which are classified in the Anophelinae and Culicinae subfamilies. Anophelinae comprises the *Anopheles, Bironella*, and *Chagasia* genera, while Culicinae subfamily includes Aedeomyiini, Aedini, Culicini, Cuisetini, Ficalbiini, Hoggesiini, Mansoniini, Orthopodomyiini, Sabethini, Toxorhynchitini, and Uranotaeniini tribes and comprises 110 genera^(1)^. About 150 mosquito species are vectors of human viral diseases. The relationship between the different taxonomic groups of viruses and mosquitoes is ancestral, with a recent description of the huge diversity of insect-specific viruses that may constitute the source of future, emerging viral diseases through species jumping from enzootic to epizootic cycles^(2)^.

Classical taxonomy has allowed us to obtain information about mosquito species’ morphology, forming groups that are not always monophyletic or similar in their distribution in ecosystems^(3)^. Additionally, classical taxonomy defines morphological characters used in dichotomic keys for species identification. However, as the knowledge of biological species has become more detailed, morphological characters may be limited or confusing and are insufficient for achieving the correct classification^(4)^. The presence of cryptic species and species complexes cannot be separated by morphological characters^(5)^ which also contributes to the problem. Molecular taxonomy is a complementary strategy to determine the relationship between species that have been difficult to determine via morphological characters, development states, and sexual dimorphism^(6)^. DNA barcoding was established in 1993 as a strategy for unifying the use of molecular markers for species identification and taxonomic allocation through phylogenetic inference based on genetic variability^(7)^. The cytochrome oxidase c subunit I (COI) gene has been widely used for molecular identification and, together with classical taxonomy, is a powerful tool for mosquito species demarcation^(8)^.

In Colombia, numerous studies have used DNA barcoding with the COI, internal transcribed spacer 2 (ITS2), and 16S subunit genes to identify *Aedes*, *Anopheles*, and *Culex* mosquitoes^(9–11)^. In the Sierra Nevada de Santa Marta (SNSM) rainforest, mosquitoes involved in arbovirus transmission have not been described in detail^(12,13)^ and there are no reports that include molecular taxonomy. In the present study, using DNA barcoding and classical taxonomy, we investigated the diversity forest mosquito species in this region of the country. The precise identification of mosquito species is essential to determine the real and potential risk of arbovirus or arthropod-borne parasite transmission and implement vector control programs.

## Materials and methods

### Study area

Mosquito specimens were collected in the SNSM foothills in Guachaca, which is in the sylvatic area of Quebrada Valencia-La Piedra (11°14’22.6” N, 73°47’58.3” W) at an altitude of 80 meters above sea level (Figure 1). The region is characterized by bimodal type rains due to the variation in precipitation, which decreases in the southeastern slope and increases in the northern slope, causing a hydrographic system to be formed. Temperatures during the year can range from 28°C to less than 0°C in the highest altitude areas, and there is a distribution of eight biomes along the SNSM depending on altitude, climatic, geographical, and physicochemical conditions^(14,15)^.

**Figure 1.**
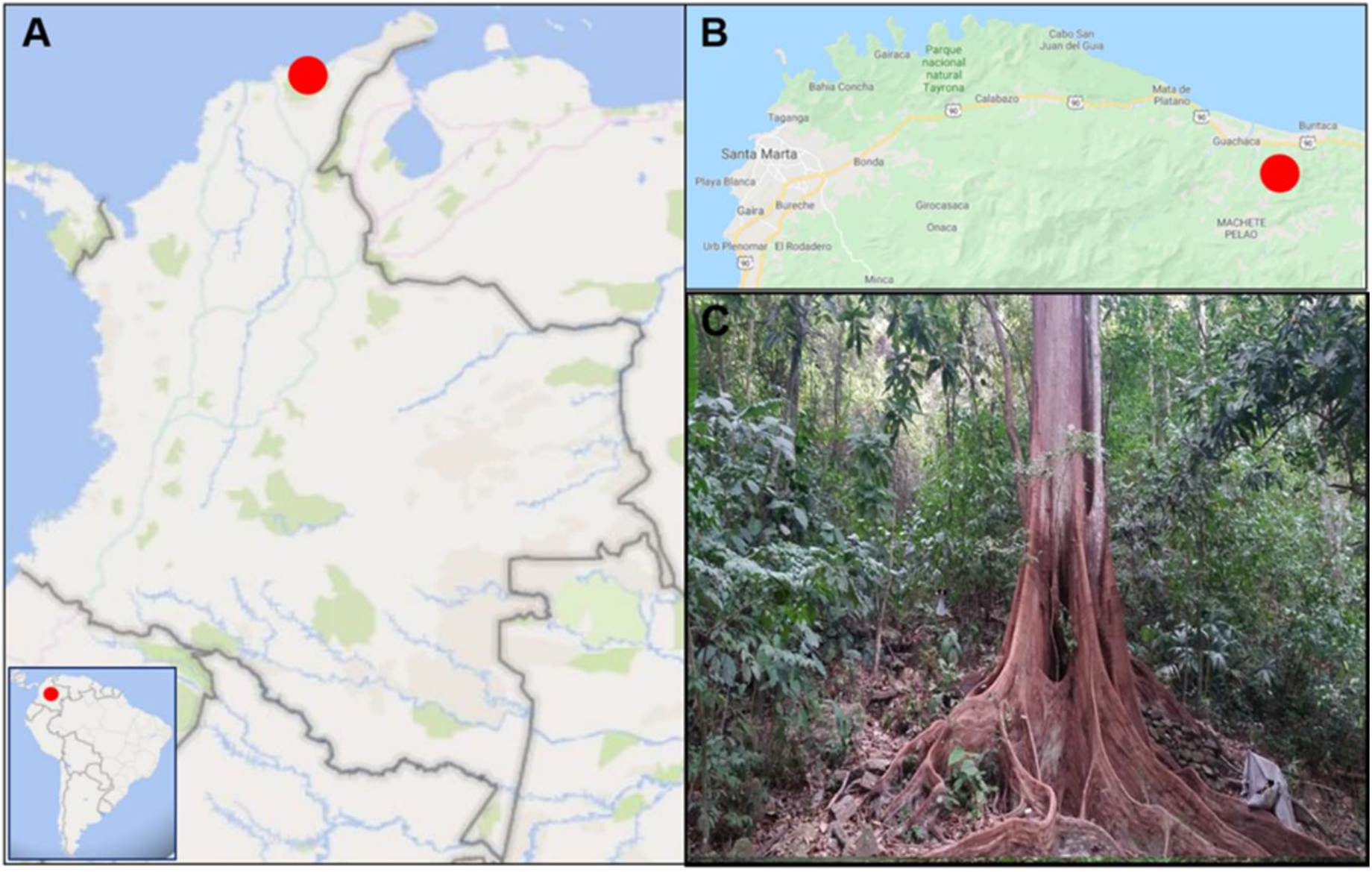
Tropical rainforest in Sierra Nevada de Santa Marta, Colombia: (A) Location of the SNSM rainforest area, (B) Guachaca locality, and (C) Quebrada Valencia-La Piedra forest.

### Mosquito collections and identification

Mosquitoes were collected during two field expeditions per month between August and December 2018, the season with the highest rainfall. Linear transects with a distance of 2 km were delimited inside the sampling area. The collection methods used were manual capture with entomological nets and aspirators. The capture was made between 07:00 and 12:00 and 14:00 and 16:00. The collected mosquitoes were transported in containers to the entomology laboratory at the Tropic Health Research Center (CIST) and sacrificed with ethyl ether. Subsequently, mosquitoes were identified using dichotomic keys^(16–19)^, and the code CIST#### was used to identify and deposit specimens into the entomological collection. Finally, the specimens were stored in vials with 1 ml absolute ethanol (96%) for subsequent molecular characterization of female mosquitoes.

### Molecular analysis

#### DNA extraction

Through the use of sterile tweezers, female legs were removed. Homogenization was performed with zirconium beads, and DNA extraction was carried out using the DNeasy Blood & Tissue kit (Qiagen, Hilden, Germany, DE).

#### COI gene amplification

The standard 658 base pairs (bp) barcode of the mitochondrial *COI* gene was amplified using the primers LCO1490 and HCO2198^(20)^. The reactions were carried out in a final volume of 25 μl. The reaction mixture included 5 μl of extracted DNA, 0.4 µM of each primer, 1.25 U *GoTaq DNA Ploymerase* (Promega), 0.2 mM of each dNTP, 1X buffer with 1.5 mM MgCl_2_, and nuclease free water. The PCR cycle conditions consisted of an initial denaturation step at 94°C for 10 minutes; followed by 35 cycles at 95°C for 60 s for denaturation, 50°C of 60 s for oligonucleotide hybridization, and 72°C of 60 s for extension; and then a final extension at 72°C of 5 min. An aliquot of each PCR product was used to visualize the expected amplicon by DNA gel electrophoresis, and the remaining volume was purified using the ExoSAP-IT^TM^ PCR Product Cleanup Reagent enzyme (Thermo Fisher Scientific Inc, catalog no. 78201.1.ML).

#### Sequencing

The purified PCR products were sequenced via Sanger sequencing. Consensus sequences were obtained by editing in DNASTAR Lasergene SeqMan Pro V 7.1.0 software^(21)^. The sequences were compared with those deposited in the BOLD Systems (boldsystems.org) and GenBank (blast.ncbi.nlm.nih.gov) databases. A matrix with the nucleotide sequences was subsequently created and aligned using the ClustalW tool^(22)^.

#### Phylogenetic analysis

The sequences did not show insertions/deletions (indels); therefore, no gap treatment was performed. The nucleotide substitution model was estimated using jModelTest^(23)^, and subsequently, the phylogenetic inference was made using the Neighbor-Joining and the Kimura 2 parameter models (K2P), using Mega 6.0 software^(24)^. The value of 2% was used as the genetic variation criterion for the COI gene^(8)^. To access the support of the phylogenetic tree topology, a resampling corresponding to 1000 bootstrap replicates was performed. The consensus tree was visualized and edited in the MEGA 6.0 software.

## Results

During our study, we collected and taxonomically identified 123 mosquitoes. The following genera were identified: *Aedes* (n = 2), *Anopheles* (n = 4), *Johnbelikinia* (n = 72), *Psorophora* (n = 25), *Sabethes* (n = 4), *Trichoprosopon* (n = 1), and *Wyeomyia* (n = 15). Seven species were identified via classical taxonomy and molecular analyses: *Aedes* (*Ochlerotatus*) *serratus* (Theobald, 1901) (Figure 2A), *Jonhbelkinia ulopus* (Dyar & Knab, 1906) (Figure 2D and E), *Psorophora* (*Janthinosoma*) *ferox* (Von Humboldt, 1819) (Figure 2B), *Sabethes* (*Sabethes*) *cyaneus* (Fabricius, 1805), *Wyeomyia* (*Triamyia*) *aporonoma* (Dyar & Knab, 1906), *Wyeomyia* (*Decamyia*) *pseudopecten* (Dyar & Knab, 1906) (Figure 2C), *Wyeomyia* (*Decamyia*) *ulocoma* (Theoblad, 1903) and it was not possible to identify the Sabethes sp. mosquito using classical taxonomy since some of structures were in poor condition. These species have been recorded in various areas of Colombia, they are described in supplementary table lV (Table SlV).

**Figure 2.**
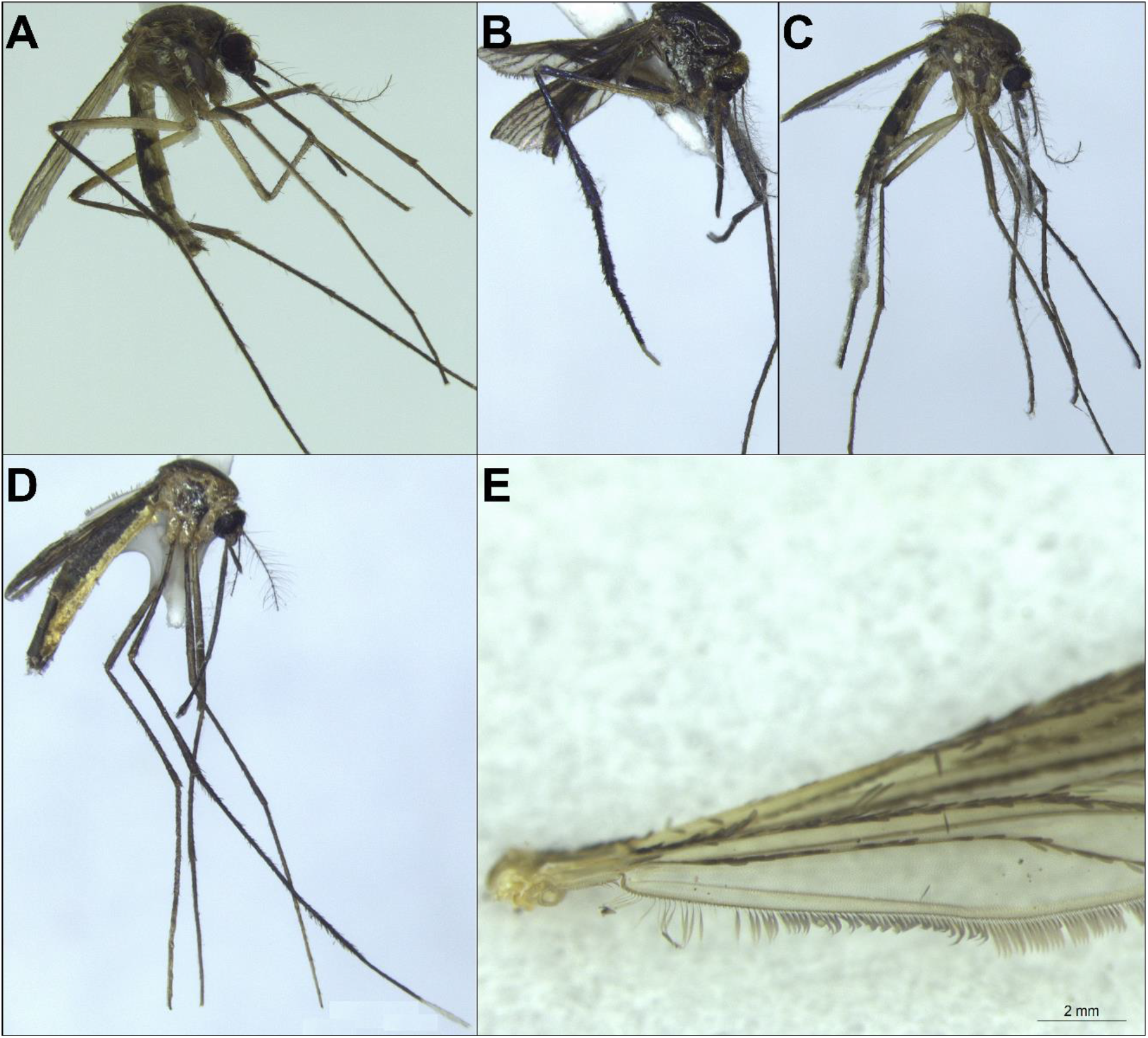
Female mosquito body of representative species. (A) *Ae. serratus*, (B) *Ps. ferox*, (C) *Wy. pseudopecten*, (D) *Jo. ulopus*, and (E) Adult stage taxonomic key of *Jo. ulopus*: upper calypter is usually without setae and rarely with 1 seta.

Seventeen sequences of the *COI* gene (658 bp) were obtained for all mosquitoes in this study (Table Sl). Eight species were confirmed through the BOLD and GenBank databases, with an identity of 90.81% to 99.61%. Only *Ae. serratus* and *Ps. ferox* had more than 15 sequences in open-access databases, and *Jo. ulopus*, *Sa. cyaneus*, *Wy. aporonoma*, *Wy. pseudopecten*, and *Wy. ulocoma* species had less than 15 accessible sequences. The *Sabethes* species was not confirmed through the databases, and its identification was ambiguous; using BOLD, an identity value <88.79% was observed in *Wy. (Wyeomyia) abebela* (Dyar & Knab, 1908), *Wy. (Wyeomyia) guatemala* (Dyar & Knab, 1906), and *Sa. chloropterus*. These sequences are not publicly accessible. In GenBank, an identity value <88.43% was observed in *Sa. (Peytonulus) hadrognatus* (Harbach, 1995) and *Wy. (Dendromyia) ypsipola* (Dyar, 1922) (Table SII).

Based on phylogenetic reconstruction, identification at the level of the Aedini, Culicini and Sabethini tribes was observed (Figure 3). Monophyletic groups were also observed. *Sabethes* sp. was located between *Johnbelikinia* and *Wyeomyia*, and a sequence of *Wyeomyia* (*Dendromyia*) *luteoventralis* (Theobald, 1901) in Colombia that does not belong to this study was located in the *Wy. Pseudopecten* group (Figure 3). Due to these two results, phylogenetic reconstruction was performed for *Aedes*, *Psorophora*, *Johnbelikinia*, *Sabethes*, and *Wyeomyia* (Figure S1-5). Species level identification was confirmed for *Ae. serratus* (Figure S1), *Ps. ferox* (Figure S2), *Jo. ulopus* (Figure S3), *Sa. cyaneus* (Figure S4), *Wy. aporonoma*, *Wy. pseudopecten*, *Wy. ulocoma*, and *Wy. luteoventralis* (Figure S5), but such confirmation was not achieved for *Sabethes* sp. since the support on the tree was <50% (Figure S4).

**Figure 3.**
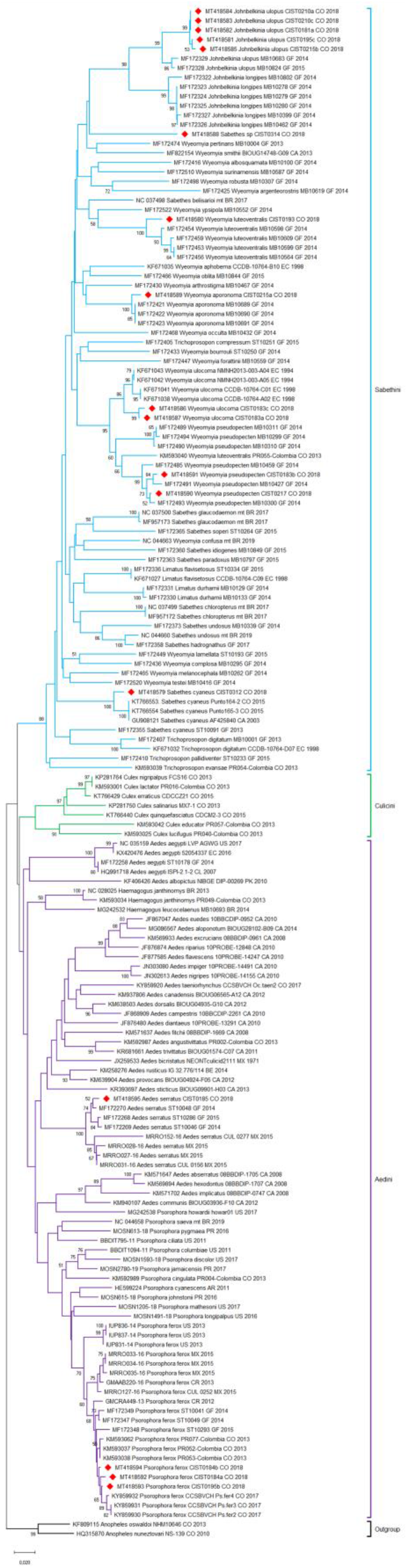
Phylogenetic reconstruction of Culicidae mosquetoes, by the NJ method, of the COI gene (353 bp). The best nucleotide substitution model was K2P, with 1000 bootstrap replicates. Sequences in the present study are highlighted with red rhombuses. An external group included *An.* (*Nyssorhynchus*) *oswaldoi* (Peryassú, 1922) and *An.* (*Nyssorhynchus*) *nuneztovari* (Gabaldón, 1940).

The intra-species genetic variation was <2% for *Wy. aporonoma* and >2% for the other species in this study (Table l). The intra-species genetic variation was <2% for *Ps. ferox* and >2% for *Sa. cyaneus* and *Wy. luteoventralis*. *Ae. serratus* reported in French Guyana and *Ps. ferox* reported in México, Costa Rica, and French Guyana were compared to the corresponding species collected and identified in this study (Tables Slll). *Sabethes* sp. was not related to species previously reported in databases and it presented with an inter-species genetic variation of >9% with the closest species (Table SIII).

**Table l.** Intra-species genetic variability for eight mosquito species. The position in the reference sequence is the length of the matrix sequences used in the phylogenetic reconstruction of Figure S1. The distance-p and the distances using the evolutionary model K2P were calculated for all sequences of the species used in this study and the species from GenBank and BOLD accessed in October 2019.

| Species | RefSeq reference sequence (GenBank access)<br>∞ | Size (bp) position in RefSeq | Average intra-species distance (distance-p) * | Average intra-species distance (K2P) + |
| --- | --- | --- | --- | --- |
| *Ae. serratus* | *Ae. aegypti* (NC_035159) | 16790: 1340-1998 | 0.0443 (4.43%) | 0.0489 (4.89%) |
| *Ps. ferox* | *Ps. saeva* (NC_044658) | 15049: 1497-2155 | 0.0211 (2.11%) | 0.0219 (2.19%) |
| *Jo. ulopus* <sup>π</sup> | *Sa. chloropterus* (NC_037499) | 15609: 1929-2587 | 0.0216 (2.16%) | 0.0227 (2.27%) |
|  | *Sa. glaucodaemon* (NC_037500) | 15620: 1930-2588 |  |  |
|  | *Sa. belisarioi* (NC_037498) | 15911: 1940-2598 |  |  |
|  | *Sa. undosus* (NC_044660) | 15334: 1518-2176 |  |  |
| *Sa. cyaneus* <sup>π</sup> | *Sa. chloropterus* | 1929-2587 | 0.0432 (4.32%) | 0.0458 (4.58%) |
|  | *Sa. Glaucodaemon* | 1930-2588 |  |  |
|  | *Sa. Belisarioi* | 1940-2598 |  |  |
|  | *Sa. Undosus* | 1518-2176 |  |  |
| *Sabethes* sp. <sup>π</sup> | *Sa. chloropterus* | 1929-2587 | - | - |
|  | *Sa. Glaucodaemon* | 1930-2588 |  |  |
|  | *Sa. Belisarioi* | 1940-2598 |  |  |
|  | *Sa. Undosus* | 1518-2176 |  |  |
| *Wy. aporonoma* | *Wy. confusa* (NC_044663) | 15426: 1515-2173 | 0.0070 (0.7%) | 0.0071 (0.71%) |
| *Wy. pseudopecten* | *Wy. confusa* | 1515-2173 | 0.0374 (3.74%) | 0.0398 (3.98%) |
| *Wy. ulocoma* | *Wy. confusa* | 1515-2173 | 0.0256 (2.56%) | 0.0269 (2.69%) |
| *Wy. luteoventralis* | *Wy. confusa* | 1515-2173 | 0.0511 (5.11%) | 0.0547 (5.47%) |

\* Number of nucleotide changes between pairs of sequences per 100 nucleotides.
+ Number of nucleotide changes (1 or more) between pairs of sequences per 100 nucleotides adjusted to the K2P evolutionary model, taking more frequent transitions than transversions.
∞ Reference sequences belonging to each genus were used since no complete mitochondrial DNA reference sequence was available in either BOLD or GenBank for the species in this study.
π *Johnbelkinia* does not have a reference sequence; therefore, the position was observed using the same sequences as *Sabethes*.

## Discussion

The present study allowed us to dissect mosquito diversity in the SNSM rainforest and inventory potential arboviral vectors in this area of the country. Our study is the first to report genetic variability of *Ae. serratus*, *Jo. ulopus*, *Wy. aporonoma*, *Wy. pseudopecten*, and *Wy. ulocoma* in the SNSM rainforest.

Sequences that lead to the identification of *Aedes serratus* and *Ochlerotatus serratus* as the same species are found in databases such as BOLD Systems and GenBank, erroneous descriptions, such as “*Ochlerotatus serratus*”, in databases creates confusion. Taxonomic reclassification has been proposed for the Aedini tribe; however, a recent analysis supported using the traditional classification^(25)^. In addition, there are difficulties in classifying the *Psorophora* genus due to similar male genitalia morphology among species^(26)^.

For *Jo. ulopus* and *Jo. longipes*, adult, larval, and pupal morphology is known, and in this study, some specimens were initially classified as *Jo. longipes* using keys that did not have the new *Johnbelkinia* classification genus, such as the keys of Cova García, 1966 and 1974, and Lane, 1953, which taxonomically classified the morphotype similar to that found for *Trichoprosopon*, the key described by Zavortink, 1979. There was a reclassification for *Trichoprosopon* and *Johnbelkinia*, via the morphological characteristic of the absence of sows in the calypter and the iridescent yellow-green colors on the mosquito scutal. These revisions allowed for the identification of *Jo. ulopus* in this study.

The Sabethini tribe has been classified as a monophyletic group using morphological characters, but when trying to classify at the genus level, there are difficulties at the morphological and phylogenetic levels since genera such as *Runchomyia*, *Tripteroides*, and *Wyeomyia*^(27,28)^ and the subgenus *Dendromyia*^(29)^ have inconsistencies, and their evolutionary relationships have not been elucidated. In addition, *Wy. compta* (Senevet & Abonnenc, 1939) and *Wy. argenteorostris* (Bonne-Wepster & Bonne, 1920) are the same species that were initially named as two different species due to classification errors by classical taxonomy^(30)^. It is essential that the taxonomic keys used are specified, and the taxonomic classification updates are taken into account, with the largest number of morphological characters analyzed.

Studies with *Ae. aegypti* showed a close phylogenetic relationship with *Hg. (Haemagogus) equinus* (Theobald. 1903)^(31)^, with some *Psorophora* members also closely related to *Aedes*^(32)^. *Ae. serratus* showed a close relationship with sequences from French Guyana and a more distant relationship with sequences from Mexico (Figure 3, S1 y Table SIlI). Currently, the circulation of this species is in the jungle. Although larvae and adults have been found at an intra and extra-domicile level in low abundances, it was not significant to define mosquito circulation in urban areas^(18)^.

The wide circulation of *Ps. ferox* has allowed the establishment of populations that begin to have morphological differences with intra-species variability in the egg and exochorion^(33)^. Genetic variability was observed in South, Central, and North America (Figure S2 and Table SIII), which suggests that conducting morphological studies in these geographical regions to identify changes in the life cycle stages may be warranted.

*Wy. aporonoma* could have a wide distribution and circulation in the mountain ranges and tropical forests present in the South American countries, although there are only reports in French Guiana and Colombia at this time ^(34)^.

The species reported as *Wy. luteoventralis* in the department of Antioquia^(10)^ was identified in the BOLD database with an identity of 98.16% and an identify of 95.74% in GenBank with *Wy. pseudopecten.* It had a closer phylogenetic relationship with the *Wy. pseudopecten* group (Figure 3 and S5), and the intra-species genetic variation with the mosquito in this study was >18% (Table SIII), which suggested an error in the taxonomic and molecular identification for this species. Therefore, the sequence analyzed would not correspond to any previously mentioned species.

All species initially characterized in this study may be arbovirus vectors, which has public health implications regarding established arboviral transmission via sylvatic and urban cycles, as well as sylvatic circulation and maintenance of arboviruses of unknown etiology that could spill over into human populations. Additionally, *Ae. serratus*, *Sa. chloropterus*, and *Ps. ferox* may have been bridge vectors that led to the establishment of the YFV sylvatic cycle^(12)^. In the future, these mosquitoes may also serve as vectors to maintain the sylvatic cycle for arboviruses such as DENV, CHIKV, and ZIKV ^(35)^.

The biogeographic characteristics of the SNSM rainforest make it a rich area of speciation due to its isolation. In addition, it is close to the Serranía del Perijá and other natural parks in the region that have been classified as speciation zones^(36)^ due to the high diversity of fauna^(37)^ and flora^(38)^. This study allowed us to corroborate the complementarity that exists between classical and molecular taxonomy. Further molecular taxonomy studies are required due to limitations of classical taxonomic keys. The keys used may be outdated regarding morphological characters of the species described; molecular identification tools are not confined by this limitation and have allowed for the expansive characterization of jungle mosquitoes via genes^(39)^ or complete mitochondrial genomes to identify species and reconstruct phylogenies^(40)^.

Possible endemism in the region reinforces the importance of developing regional DNA barcoding libraries for molecular species identification. In this study, 90% of species were identified molecularly, and 10% were identified via classical taxonomy. These differences could be explained due to sequence divergence in species and limitations of classical taxonomy.

## Acknowledgments

This study was financed and supported by the Ministry of Science, Technology, and Innovation in Health (Minciencias) Colombia, project 210477757671 (Virologic Expedition in Representative Ecosystems of Colombia: Tropical Rainforest of the Sierra Nevada de Santa Marta) to JAU-C. We thank entomologist Victor Alberto Olano from the Health and Environment group of the Universidad El Bosque for his support with species identification via classical taxonomy. We also thank technicians Adalberto Duica and Juan Domínguez (Department of Health, Santa Maria District) for field work assistance.

## Authors contributions

All authors participated equally in the work.

