## Supplementary material for "Molecular characterization of mosquitoes (Diptera: Culicidae) from the tropical rainforest of Sierra Nevada de Santa Marta, Colombia": Fig S1-S5, tables S1-S4

**Table SI.** List mosquito species *COI* sequences.

| Species | ID GenBank | ID BOLD Systems | Code CIST |
|---|---|---|---|
| *Ae. serratus* | MT418595 | CIST001-20 | 0185 |
| *Ps. ferox* | MT418592 | CIST004-20 | 0184a |
| | MT418593 | CIST003-20 | 0185b |
| | MT418594 | CIST002-20 | 0184b |
| *Jo. ulopus* | MT418581 | CIST015-20 | 0195 |
| | MT418582 | CIST014-20 | 0181a |
| | MT418583 | CIST013-20 | 0210c |
| | MT418584 | CIST012-20 | 0210a |
| | MT418585 | CIST011-20 | 0215b |
| *Sa. cyaneus* | MT418579 | CIST017-20 | 0312 |
| *Sabethes* sp. | MT418588 | CIST008-20 | 0314 |
| *Wy. aporonoma* | MT418589 | CIST007-20 | 0215a |
| *Wy. pseudopecten* | MT418590 | CIST006-20 | 0217 |
| | MT418591 | CIST005-20 | 0183b |
| *Wy. ulocoma* | MT418586 | CIST010-20 | 0183c |
| | MT418587 | CIST009-20 | 0183a |
| *Wy. luteoventralis* | MT418580 | CIST016-20 | 0193 |

**Table SII.** Molecular identification of species via database searching and availability of COI sequences, showing the first 99 results. Data accessed in October 2019.

| Species | Number of samples | BIN Code | Similarity (%) BIN* | GenBank Code | Similarity (%) GenBank | Notes |
|---|---|---|---|---|---|---|
| *Ae. serratus* | 1 | BOLD:AAN3110[+]<br><br><br>BOLD:ACN3711<br><br>BOLD:ADQ7713<br>BOLD:ADG1168<br>BOLD:ADR0304 | 99.78<br>98.86<br>98.48<br>97.92-<br>97.54<br>97.14<br>-<br>- | MF172270[+]<br>MF172268[+]<br>MF172269[+] | 99.06<br>98.87<br>98.49 | 13 private sequences in the BOLD |
| **Total free Access sequences** | | 17 | | 3 | | |
| *Ps. ferox* | 3 | BOLD:AAO0580[+]<br><br>BOLD:ABZ5766<br><br>BOLD:ACC4707<br>BOLD:ADR1076<br>BOLD:ADR1503 | 99.61 -<br>98.65<br>98.46 -<br>98.07<br>97.05<br>-<br>- | MG242536[+]<br>KM452775[+] | 99.06<br>99.12 | 28 private sequences in the BOLD |
| **Total free Access sequences** | | 181 | | 24 | | |
| *Jo. ulopus* | 5 | BOLD:ACZ4300[+] | 97.22 -<br>95.83 | MF172329[+]<br>MF172328[+] | 96.30<br>95.83 | 3 private sequences in the BOLD |
| **Total free Access sequences** | | 2 | | 2 | | |
| *Sa. cyaneus* | 1 | BOLD:AAX9629[+] | 98 | GU908121[+]<br>KT766553[+] | 97.59<br>- | |

| | | | | | | |
|---|---|---|---|---|---|---|
| | | <br>BOLD:ACZ3883[+] | <br>90.89 | KT766554[+]<br>MF172355[+] | -<br>90.81 | |
| **Total free Access sequences** | | 4 | | 5 | | |
| *Sabethes sp* | 1 | - | - | - | - | |
| **Total free Access sequences** | | - | - | - | - | |
| *Wy. aporonoma* | 1 | BOLD:ACZ4142[+] | 98.6 | MF172423[+]<br>MF172422[+]<br>MF172421[+] | 98.60 | 19 private sequences in the BOLD |
| **Total free Access sequences** | | 3 | | 3 | | |
| *Wy. pseudopecten* | 2 | BOLD:AAG3839[+]<br>BOLD:ADZ5713[+]<br><br>BOLD:ADZ5714[+]<br>BOLD:ACU5366[+] | 99.84<br>97.98<br><br>97.83<br>94.88 | MF172493[+]<br>MF172487[+]<br>MF172485[+]<br>MF172491[+]<br>KM593040[+] | 99.53<br>97.98<br>97.83<br>97.83<br>94.88 | 5 private sequences in the BOLD |
| **Total free Access sequences** | | 10 | | 10 | | |
| *Wy. ulocoma* | 2 | BOLD:AAG3840[+] | 96.28 - 95.66 | KF671038[+] | 96.28 – 95.66 | 1 private sequences in the BOLD |
| **Total free Access sequences** | | 8 | | 8 | | |
| *Wy. luteoventralis* | 1 | BOLD:ACZ3898[+]<br>BOLD:ACU5366[+] | 97.02<br>- | MF172452[+]<br>MK593949[+] | 97.35<br>- | |
| **Total free Access sequences** | | 10 | | 10 | | |

* Barcode Index Numbers (BINs)
+ Report of a sequence in BOLD and GenBank

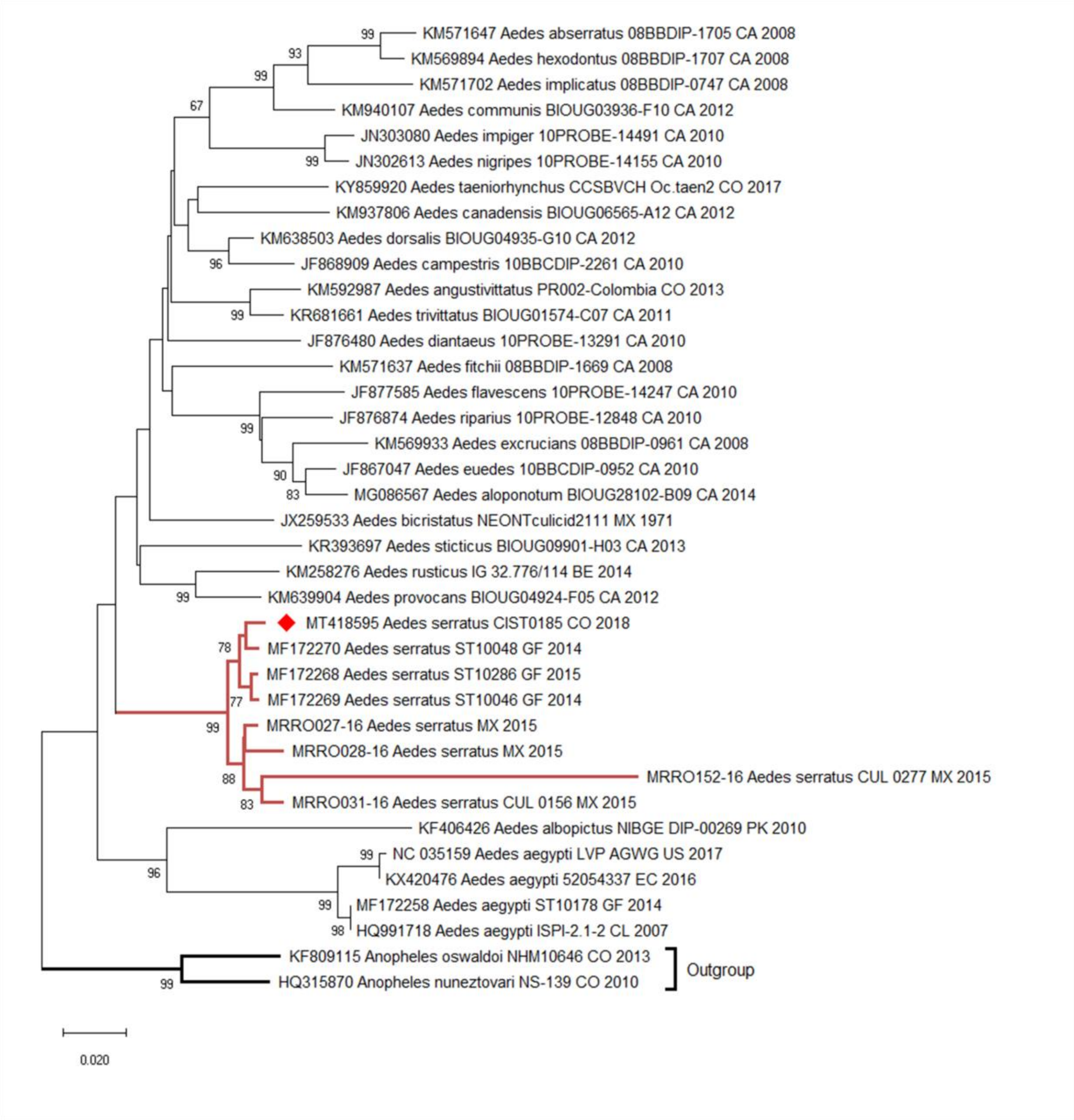


**Figure S1.** Phylogenetic reconstruction of *Aedes* (530 bp) by NJ for the COI gene. The best nucleotide substitution model was K2P, 1000 bootstrap replicates. Sequences in the present study are identified with red rhombuses. External groups: *An. oswaldoi* y *An. nuneztovari*.

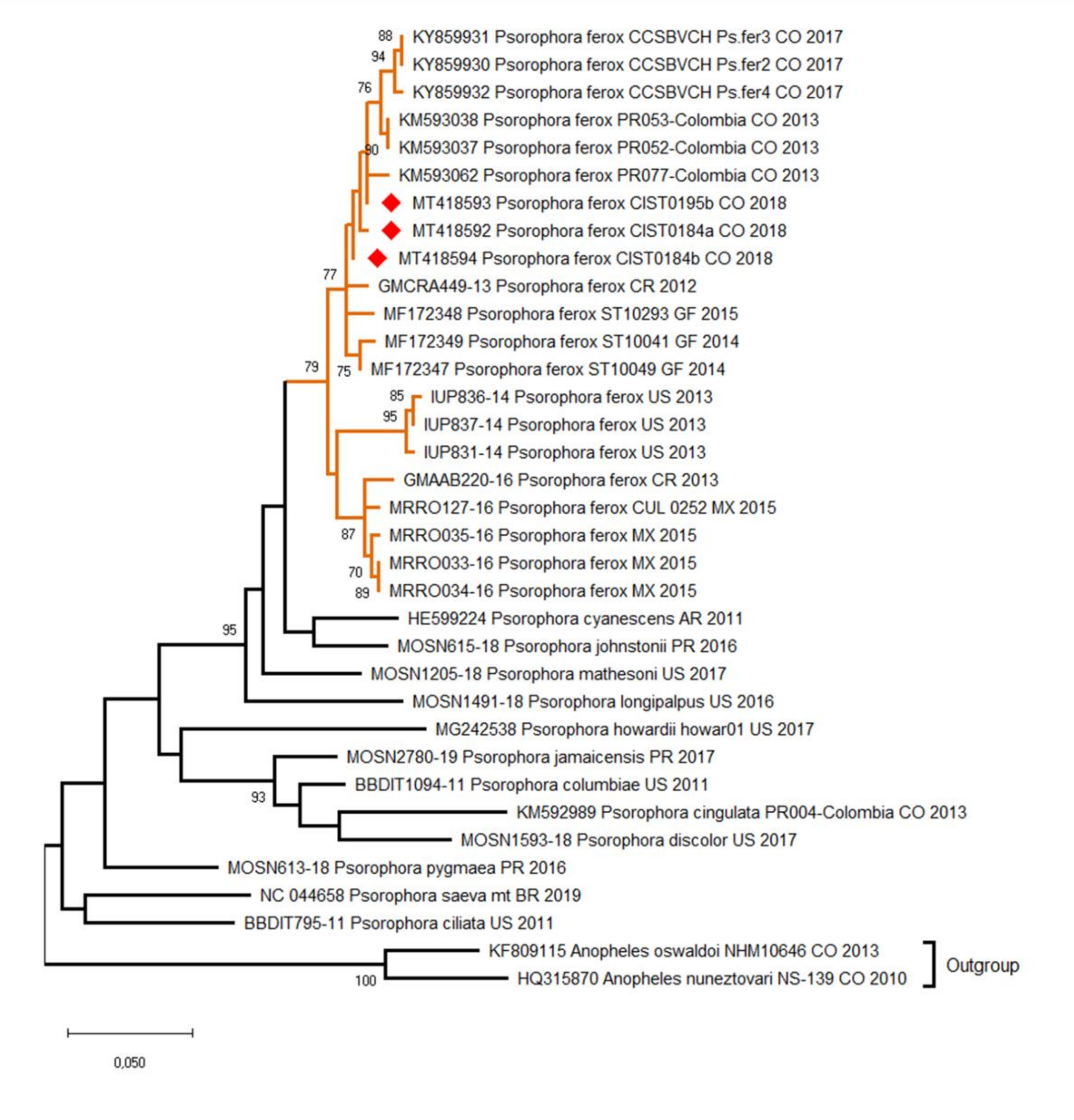


**Figure S2.** Phylogenetic reconstruction of *Psorophora* (530 bp) by NJ for the COI gene. The best nucleotide substitution model was K2P, 1000 bootstrap replicates. Sequences in the present study are identified with red rhombuses. External groups: *An. oswaldoi* y *An. nuneztovari*.

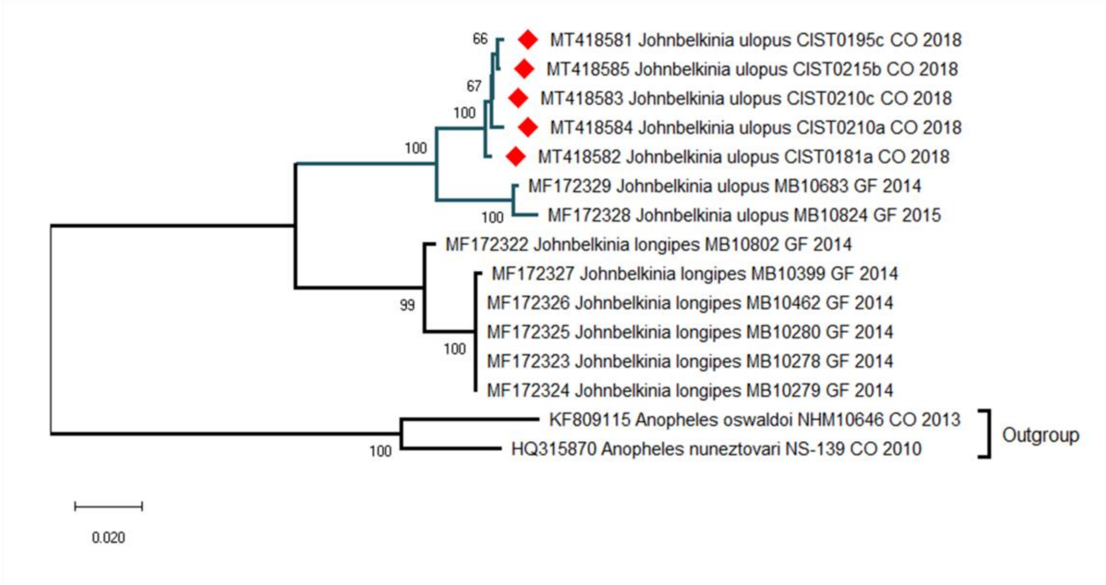


**Figure S3.** Phylogenetic reconstruction of *Johnbelikinia* (648 bp) by NJ for the COI gene. The best nucleotide substitution model was K2P, 1000 bootstrap replicates. Sequences in the present study are identified with red rhombuses. External groups: *An. oswaldoi* y *An. nuneztovari*.

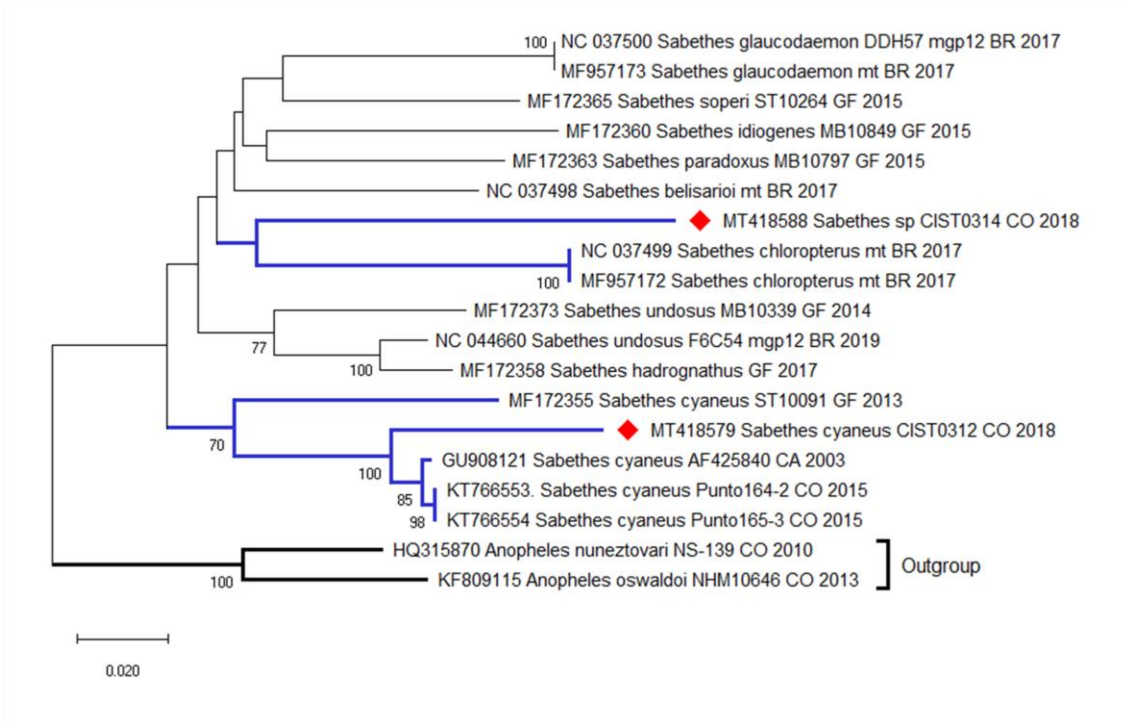


**Figure S4.** Phylogenetic reconstruction of *Sabethes* (456 bp) by NJ for the COI gene. The best nucleotide substitution model was K2P, 1000 bootstrap replicates. Sequences in the present study are identified with red rhombuses. External groups: *An. oswaldoi* y *An. nuneztovari*.

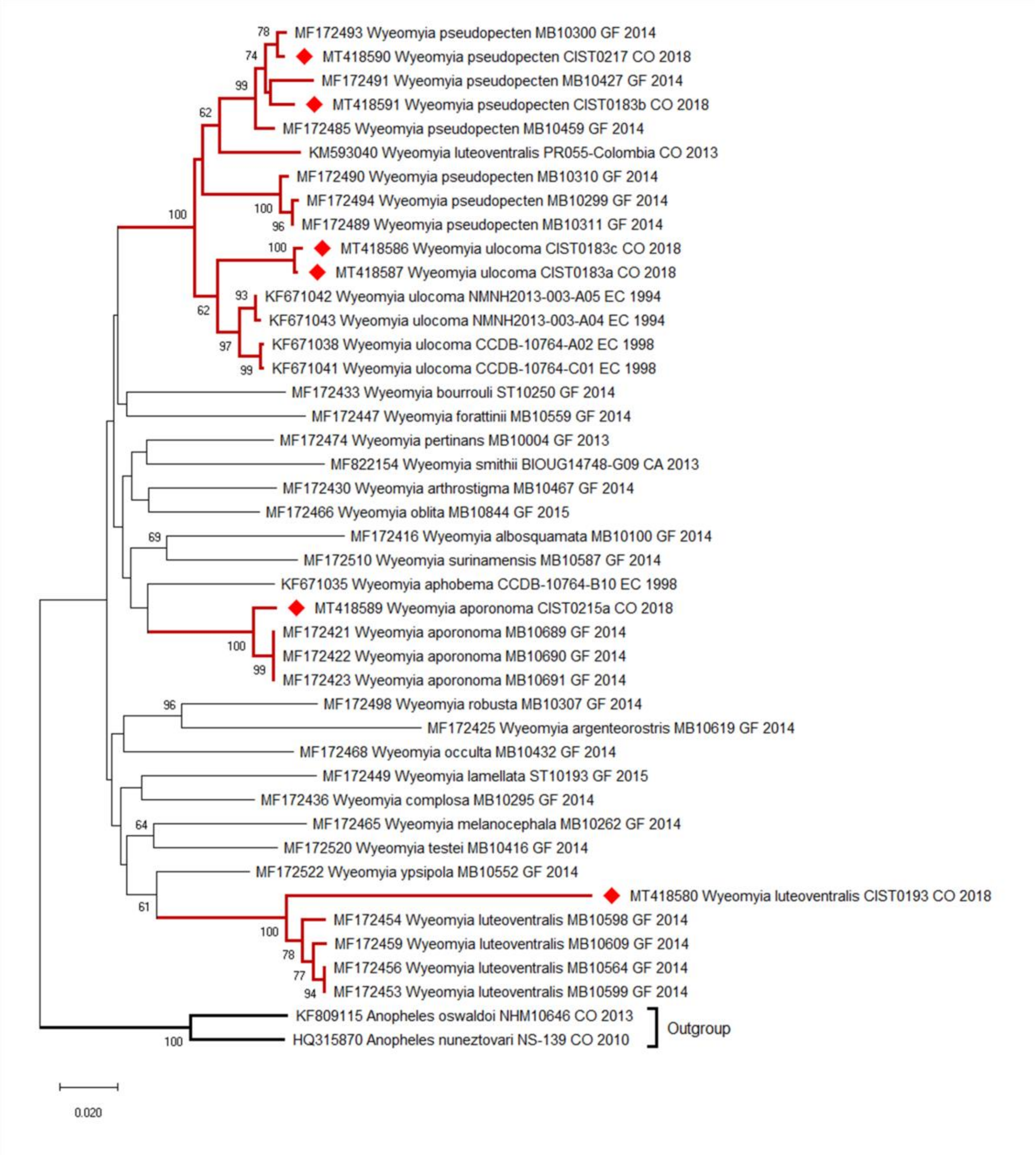


**Figure S5.** Phylogenetic reconstruction of *Wyeomyia* (645 bp) by NJ for the COI gene. The best nucleotide substitution model was K2P, 1000 bootstrap replicates. Sequences in the present study are identified with red rhombuses. External groups: *An. oswaldoi* y *An. nuneztovari*.

**Table SIII.** Intra- and inter-species genetic variability for the species in this study. The distance-p and the distance using the evolutionary model K2P were calculated by grouping countries and described using two-digit codes.

| Species | Codes | Average of distance (distance-p) | Average of distance (K2P) |
|---|---|---|---|
| *Ae. serratus* | Ae1:CO*-MX | 6.1 | 6.75 |
| *Ae. serratus* | Ae2:CO*-GF | 1.0 | 1.05 |
| *Ps. ferox* | Ps1:CO*-CO$^{\psi}$ | 0.85 | 0.86 |
| *Ps. ferox* | Ps2:CO*-CO+ | 0.65 | 0.67 |
| *Ps. ferox* | Ps3:CO*-CO~ | 0.84 | 0.86 |
| *Ps. ferox* | Ps4:CO*-CR | 1.3 | 1.33 |
| *Ps. ferox* | Ps5:CO*-US | 2.11 | 2.20 |
| *Ps. ferox* | Ps6:CO*-MX | 1.29 | 1.33 |
| *Ps. ferox* | Ps7:CO*-GF | 0.84 | 0.85 |
| *Sa. cyaneus* | Sa1:CO*-CO^ | 1.97 | 2.01 |
| *Sa. cyaneus* | Sa2:CO*-GF | 8.99 | 9.6 |
| *Sa. cyaneus* | Sa3:CO*-CA | 1.97 | 2.01 |
| *Sabethes sp* | Sa4:CO*-A | 9.21 | 10.77 |
| *Sabethes sp* | Sa5:CO*-B | 13.82 | 16.22 |
| *Sabethes sp* | Sa6:CO*-C | 12.62 | 14.64 |
| *Sabethes sp* | Sa7:CO*-D | 16.89 | 20.49 |
| *Wy. pseudopecten* | Wy1:CO*-E | 4.06 | 4.32 |
| *Wy. luteoventralis* | Wy2:CO*-E | 11.63 | 12.65 |

CO* Sequences obtained from this study.
CO$^{\psi}$ Colombian sequences reported in BOLD and GenBank.
CO+ Colombian sequences reported in the department of Antioquia, La Pintada municipality.
CO~ Colombian sequences reported in the department of Córdoba, Montería municipality, San Bernardo del Viento, Village Chiquin.
CO^ Colombian sequences reported in the department of Córdoba, Montería municipality, San Bernardo del Viento, La Balsa.
A. *Sa. chloropterus* sequence
B. *Sa. hadrognathus* sequence
C. *Sa. belisarioi* (Min value) sequence
D. *Sa. cyaneus* (Max value) sequence
E. *Wy. luteoventralis* sequence reported in the department of Antioquia, La Pintada municipality.

**Table SIV.** Geographical records of mosquitoes in Colombia and associated arboviruses. Data accessed in October 2019.

| Species | Department | Reference | Associated arbovirus | References |
|---|---|---|---|---|
| *Wy. aporonoma* | Valle | Heinemann & Belkin, 1978<br>Barreto *et al*, 1996 | -- | -- |
| *Wy. luteoventralis* | Antioquia | Rozo-López & Mengual, 2015 | -- | -- |
| *Jo. ulopus* | Boyacá<br>Meta<br>Nariño<br>Norte de Santander<br>Valle del cauca<br>Antioquia<br>Caldas | Suaza-Vasco *et al*, 2015 | -- | -- |
| *Wy. pseudopecten* | Valle del cauca | Suaza-Vasco *et al*, 2015 | -- | -- |
| *Wy. ulocoma* | Valle del Cauca | Suaza-Vasco *et al*, 2015 | -- | -- |
| *Sa. cyaneus* | Meta | Bates, 1945 | | |
| | Valle del Cauca | Suaza-Vasco *et al*, 2015 | | |
| | Caquetá | Molina *et al*, 2014 | | |
| | Córdoba | Hoyos-López *et al*, 2015 | | |
| *Sa. chloropterus* | Meta | Bates, 1945 | Primary vector: Yellow Fever Virus (YFV) | Galindo *et al*, 1958<br>Galindo *et al*, 1956<br>Zsemlye *et al*, 2005 |
| | Valle del Cauca | Suaza-Vasco *et al*, 2015 | | |
| | Caquetá | Molina *et al*, 2014 | | |
| | Córdoba | Hoyos-López *et al*, 2015 | | |
| *Ae. serratus* | Antioquia | López, 1990<br>Groot, 1974<br>Barreto *et al*, 1996<br>Parra-Henao & Suarez, 2012 | Secondary vector: Yellow Fever Virus (YFV) | Cardoso *et al*, 2010<br>Sick *et al*, 2019<br>Pinheiro *et al*, 2019 |
| | Meta | Antunes, 1937 | | |
| | Caquetá | Molina *et al*, 2014 | Mayaro virus (MAYV) | Muñoz *et al*, 2012 |
| | Córdoba | Heinemann & Belkin, 1978<br>Morales & Valdez, 1962 | Venezuelan Equine Encephalitis virus (EEV) | Molina *et al*, 2014 |
| | Valle del Cauca | Barreto & Lee, 1969 | | |
| | Santander | Ferro *et al*, 2008<br>Groot *et al*, 1961 | | |
| *Ps. ferox* | Antioquia | Rozo-López & Mengual, 2015<br>Hoyos-López, 2018<br>Parra-Henao & Suarez, 2012 | West Nile virus (WNV) | Christofferson *et al*, 2010 |
| | | | Eastern Equine Encephalitis (EEEV) | Navia-Gine *et al*, 2013<br>Oliver *et al*, 2018 |
| | Vichada | Maria & Castro, 2016 | St Louis Encephalitis virus (SLEV) | Beranek *et al*, 2018 |
| | Valle del Cauca | Figueroa, 1953 | Madariaga (MADV) | Lednicky *et al*, 2019 |
| | Caquetá | Molina *et al*, 2014 | Venezuelan Equine Encephalitis virus (EEV) | Molina *et al*, 2014 |
| | Guajira | Morales *et al*, 2014 | | |
| | Magdalena | Dickerman *et al*, 1987 | | |
